# Real-time, High-throughput Super-resolution Microscopy via Panoramic Integration

**DOI:** 10.1101/2025.08.07.669117

**Authors:** Kyungduck Yoon, Hansol Yoon, Kidan Tadesse, Zhi Ling, Biagio Mandracchia, Sayantan Datta, G Ozan Bozdag, Anthony J Burnetti, William C Ratcliff, Shu Jia

## Abstract

We introduce super-resolution panoramic integration (SPI), an on-the-fly microscopy technique enabling instantaneous generation of subdiffractional images concurrently with scalable, high-throughput screening. SPI leverages multifocal optical rescaling, high-content sweeping, and synchronized line-scan readout while preserving minimal post-processing and compatibility with epi-fluorescence settings. We demonstrate SPI for various subcellular and populational morphology, function, and heterogeneity. This versatile platform offers a practical pathway toward biological insights beyond traditional optical and computational constraints.

## INTRODUCTION

Modern single-cell analysis necessitates imaging technologies that combine high spatiotemporal resolution, multiparametric content, and robust throughput for scalable visualization and profiling of cell populations without compromise^1–3^. Super-resolution microscopy has overcome the physical diffraction barriers inherent in traditional optical microscopes, revealing subcellular details with unprecedented clarity^4–7^. Among major super-resolution techniques, structured illumination microscopy is particularly noteworthy for balancing resolution enhancement with moderate acquisition speeds, low phototoxicity, and compatibility with general sample preparation^8,9^. Meanwhile, the emergence of image scanning microscopy exploits a confocal variant to improve optical sectioning, signal-to-noise ratios, and acquisition speed^10,11^. Current image-scanning implementations primarily depend on complex scanning hardware, such as confocal spinning disks, galvanometric mirrors, or digital micromirror devices^12–18^, which may hinder convenient integration into commonly available microscopes. In contrast, recent refinements into multifocal scanning microscopy replace beam-scanning optics with stationary components and physical sample translation^19,20^, simplifying the instrumentation and enabling super-resolution imaging with conventional epi-fluorescence setups. However, current methods remain limited in acquiring high contents and often rely on post-processing for image reconstruction, thereby restricting their applicability for rapid, largescale screening and visualization in single-cell analysis.

Here, we present super-resolution panoramic integration microscopy (SPI), which enables the instant formation of super-resolution images in parallel with scalable, high-throughput screening of biological specimens. SPI leverages multifocal optical rescaling, high-content sample sweeping, and synchronized sensor line-scan readout while reducing post-processing and preserving conventional epi-fluorescence settings. The technique effectively achieves twofold resolution enhancement, unperturbed streaming throughput, technically unconstrained field of view (FOV), and, notably, super-resolution outputs on the fly. We validate SPI across diverse phantom and biological specimens, demonstrating rapid, population-level analysis of biological systems. Collectively, these capabilities establish SPI as a versatile platform with transformative potential for high-throughput biological discovery, offering a practical pathway for advancements in cell biology, pathology, and large-scale diagnostic applications.

## RESULTS

### Principle of SPI and system characterization

In principle, SPI streamlines optical photon reassignment, high-content sample sweeping, and in-phase time-delay integration (TDI) readout through an epi-fluorescence setup (**Fig. 1a, Fig. S1, Methods**). The SPI framework circumvents the need for complex beam-scanning schemes and extensive post-processing required in existing pixel-reassignment-based approaches (**Table S1**). The SPI system incorporates concentrically aligned microlens arrays in both the illumination and detection paths, contracting point-spread functions (PSFs) by a factor of √2, thus surpassing the diffraction limit without photon loss^21,22^ (**Supplementary Note 1, Methods**). Furthermore, SPI employs a TDI sensor that synchronizes line-scan readout with the corresponding sweeping sample motion (**Fig. 1b**), offering unique advantages over conventional frame-based sensors (**Supplementary Note 2, Table S1**). This real-time synchronization enables full-specimen capture and, notably, instant and unperturbed formation of super-resolved images with samples being continuously introduced through the FOV, eliminating delays associated with computational reconstructions (**Supplementary Note 3, Methods**). Implementing non-iterative rapid Wiener-Butterworth (WB) deconvolution^25^ provides an additional √2× enhancement in resolution, obtaining the full 2× improvement consistent with standard SIM techniques (**Fig. 1c, Fig. S2, Table S1**). It should be mentioned that WB-processing offers ~40-fold faster processing (down to 10 ms) compared to traditional Richardson-Lucy deconvolution, making it particularly advantageous for high-throughput image analysis (**Fig. S2**).

**Figure 1:**
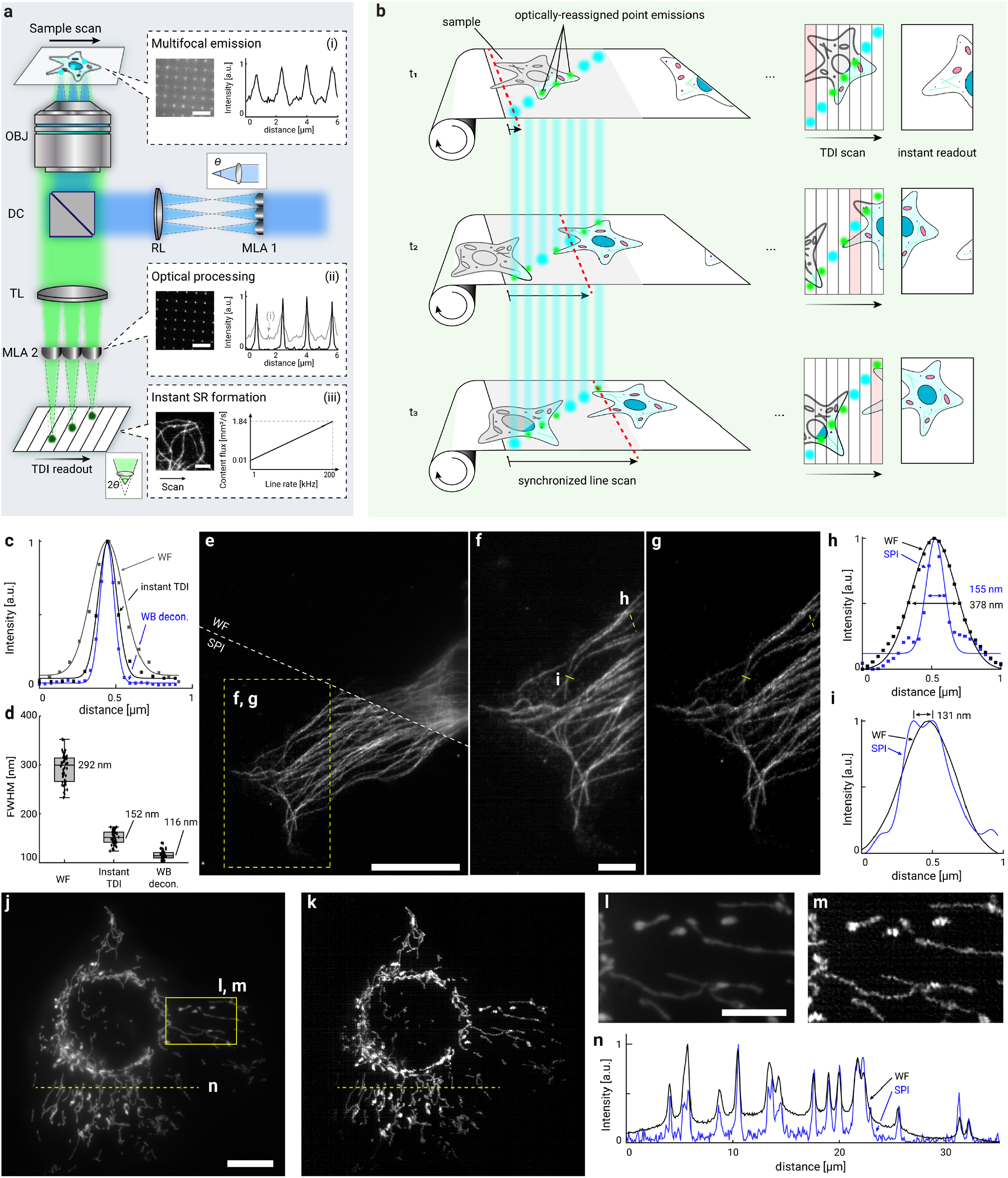
Super-resolution panoramic integration (SPI). (a) Schematic of the SPI setup. Paired microlens arrays (MLAs) enable optical photon reassignment via focal spot scaling. OBJ, objective lens; DC, dichroic cube; RL, relay lens; TL, tube lens. Insets: (i, ii) Left, fluorescent emission upon multifocal illumination (i) and after optical photon reassignment (ii); right, corresponding cross-sectional profiles over multiple emission spots, showing the PSF contraction (*θ* to 2*θ*) with enhanced sectioning through out-of-focus rejection. (iii) Left, instant super-resolution image formation from the TDI sensor; right, FOV and content flux as a function of line readout rate. (b) Illustration of the SPI workflow. Synchronous line-scan readout with sweeping sample motion across an unperturbed recording on the fly while displaying instantaneous super-resolved images. (c, d) Wide-field (WF), instant TDI readout, and WB-refined cross-sectional profiles (c) and analysis (d) of multiple fluorescent point emitters (n=50), showing the FWHM values of 292 ± 29 nm, 152 ± 13 nm, and 116 ± 9 nm, respectively. (e) WF and SPI images of *β*-tubulin in HeLa cells. (f, g) Zoomed-in WF (f) and SPI (g) images of the boxed region in (e). (h, i) Cross-sectional intensity profiles, as marked in (f, g), revealing adjacent structures separated by ~130 nm and ~2× resolution improvement. (j, k) WF (j) and SPI (k) images of GFP-tagged mitochondria in U2OS cells. (l, m) Zoomed-in WF (l) and SPI (m) images of the boxed region in (j). (n) Cross-sectional intensity profiles, as marked in (j, k), revealing enhanced optical sectioning, clarity, and resolution of mitochondrial structures using SPI. Scale bars: 3 μm (a), 20 μm (e), 5 μm (f), 10 μm (j), 5 μm (l).

Testing fluorescent point emitters, the instant TDI readout produced a narrower sub-diffraction-limited FWHM (152 ± 13 nm), which can be further refined with WB deconvolution (116 ± 9 nm) (**Fig. 1c-d, Fig. S3**). These PSF measurements contrast conventional wide-field images (292 ± 29 nm) acquired by integrating unassigned point emissions (**Fig. S4**). We further validated the performance of SPI using biological samples, including *β*-tubulin, mitochondria, and peroxisomes (**Fig. 1e-n, Fig. S3, and Fig. S5**). As seen, super-resolved SPI delineated subcellular structures with increased clarity and sectioning and exhibited a consistent two-fold resolution enhancement (~120 nm). In addition, these results across diverse biological specimens verified continuous super-resolution streaming with unperturbed high throughput (**Supplementary Videos S1-3**), acquiring up to 1.84 mm^2^/s, typically containing 5,000-10,000 cells/s (**Supplementary Note 4**).

### High throughput, super-resolution imaging of peripheral blood smear

Leveraging raster scanning, SPI effectively scales super-resolution imaging to accommodate large cellular populations and whole-slide applications. We demonstrated this capability by imaging peripheral blood smears labeled with wheat germ agglutinin (WGA) (**Fig. 2a, Methods**). Specifically, SPI continuously inspected a testing area containing over 100,000 cells with modest sweeping at 9,250 μm^2^/s across 2 mm × 2 mm, exceeding the static FOVs of conventional fluorescence microscopy techniques (**Fig. 2b, Supplementary Video S4**). Notably, SPI can acquire such an area of interest using around 60 seconds (**Supplementary Video S4)**, a throughput in line with commercial whole-slide scanners (typically 40-180 seconds for an equivalent imaging area)^26^, while delivering unprecedented sub-diffraction-limited clarity of major red blood cells (**Fig. 2c-e**). Moreover, WGA-stained images generated by SPI distinctly depicted cell membranes and cytoplasm, facilitating viable three-part differential analysis of white blood cell subtypes, such as lymphocytes, neutrophils, and monocytes (**Fig. 2f-h**). Population analysis (>80,000 cells) revealed that red blood cells (93.1%) and platelets (6.8%) comprise the majority of the blood smear. Owing to the high sensitivity of SPI, rare white blood cells were resolved into neutrophils (49.3%), lymphocytes (38.8%), and monocytes (11.9%), in close relevance to conventional hematologic quantifications^27^ (**Fig. 2i**).

**Figure 2.**
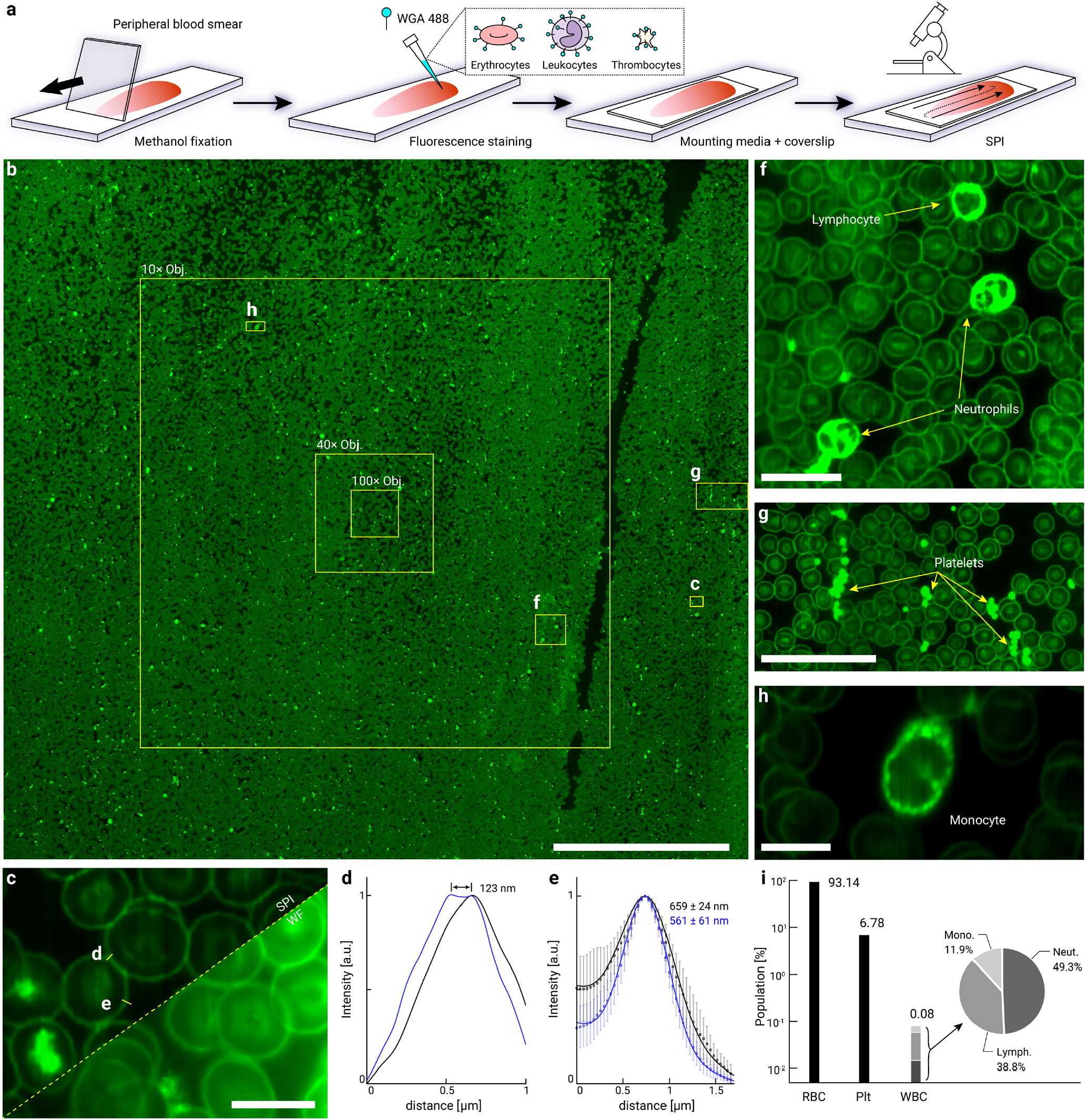
Imaging peripheral blood smear with SPI. (a) Illustration of the SPI workflow for imaging blood smear labeled for wheat germ agglutinin (WGA), staining red blood cell (RBC) membranes, white blood cell (WBC) cytoplasm, and platelets. (b) Super-resolution SPI image of raster-scanned millimeter-sized blood smear areas, in comparison with static FOVs of conventional fluorescence microscopy. (c-e) WF and SPI images of RBCs (c) and corresponding cross-sectional intensity profiles (d, e), indicating better optical sectioning and SPI resolution enhanced by over 100 nm compared to the diffraction limit. (f-g) Three-part differential analysis enabled by SPI, clearly resolving RBCs, lymphocytes, neutrophils, monocytes, and platelets. (i) Population analysis (>80,000 cells) based on the SPI results, closely relevant to standard hematology reference values of 95: 4.9: 0.1 (RBC, platelet, WBC) and 65: 30: 5 for WBC subtypes (neutrophils, lymphocytes, monocytes), respectively. Scale bars: 0.5 mm (b), 10 µm (c), 20 µm (f), 50 µm (g), 10 µm (h).

### Analyzing morphological features of snowflake yeast clusters

Next, we employed SPI to image eGFP-tagged snowflake yeast (*Saccharomyces cerevisiae*), a model system for studying the evolution of multicellularity^28–30^ (**Fig. 3a, Methods**). Snowflake yeast form through incomplete daughter cell separation due to deletion of the transcription factor *ACE2*, resulting in clusters that grow via mother-daughter cell attachment in a fractal-like branching pattern^28^. These clusters reproduce when mechanical strain from cellular packing breaks cell-cell bonds, liberating multicellular propagules^29,30^. In the Multicellularity Long Term Evolution Experiment (MuLTEE), populations that started out microscopic (~32 µm in diameter) subjected to daily settling selection for increased size evolved to be macroscopic (~mm scale^29^; **Fig. 3a, b**). This remarkable size increase occurs primarily through branch entanglement, where elongated cells wrap around one another, creating mechanically robust networks that resist fracture even when cell-cell bonds break^29^.

**Figure 3.**
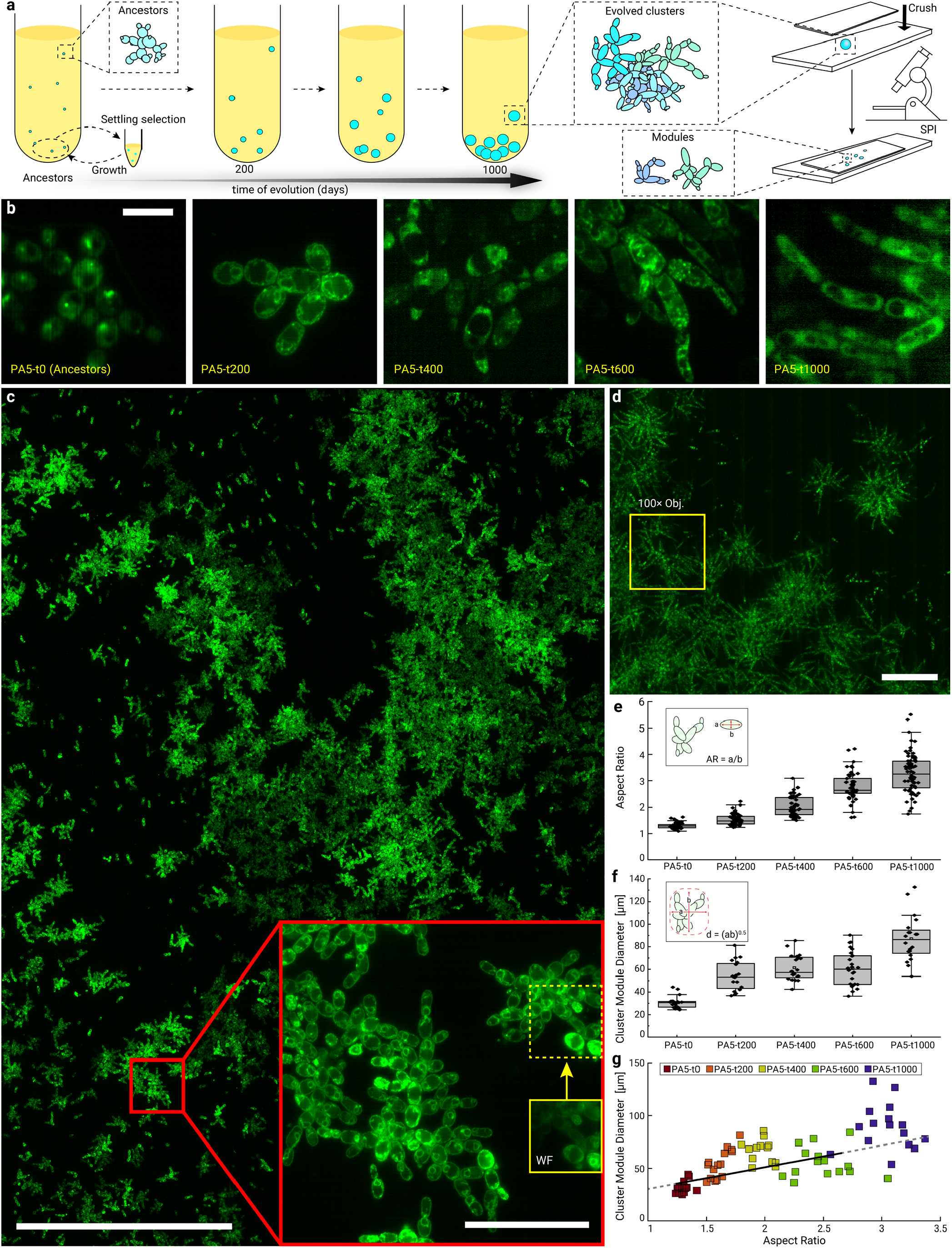
Imaging and quantification of multicellular snowflake yeast phenotype (*Saccharomyces cerevisiae*). (a) Diagram of the SPI workflow for preparing and imaging snowflake yeasts under varying evolutionary stages of anaerobic conditions with settling selection. (b) Exemplary SPI images of the ancestral (t0) and evolved PA5-t200, PA5-t400, PA5-t600, and PA5-t1000 snowflake yeast clusters, with stage-specific GFP tagging on PDC1 (ancestors), the endoplasmic reticulum (PA5-t200 and PA5-t600), and HSC82 (PA5-t400 and PA5-t1000). PA, anaerobic population. (c) Super-resolution SPI image of raster-scanned PA5-t200 snowflake yeast, revealing the finer cellular distribution of early-stage yeast clusters compared to wide-field microscopy, as shown in the zoomed-in inset images. (d) Super-resolution SPI image of raster-scanned PA5-t1000 snowflake yeast, displaying the cellular aspect ratios and density of later-stage yeast modules that emerge from flattened clusters. The boxed area indicates a static FOV of conventional fluorescence microscopy. (e, f) Quantitative analyses show that cellular aspect ratios (e) and cluster module size (f) increased over evolutionary time, indicating an increasing trend as a function of evolutionary time. (g) The relationship between cluster module diameter and cellular aspect ratio shows a linear trend during early evolution, with notable divergence at specific timepoints (t400 and t1000) where modules become more extensive than expected based on cellular aspect ratio, suggesting the co-evolution of adaptations that reduce strain accumulation or increase mechanical toughness. Scale bars: 10 µm (b), 0.5 mm (c), 50 µm (c inset), 100 µm (d).

To visualize subcellular features across evolutionary timepoints, we employed distinct fluorescent markers targeting different cellular components: PDC1 (ancestors), the endoplasmic reticulum (PA5-t200 and PA5-t600), and HSC82 (PA5-t400 and PA5-t1000) (**Fig. 3b**). The HSC82 tagging is particularly significant as its downregulation plays a key role in driving cellular elongation during multicellular evolution^31^.

When compressed under a coverslip for imaging, these macroscopic clusters disassociate into smaller “modules” that resemble the ancestor clusters. These modules have characteristic size distributions - a property not yet systematically explored in the MuLTEE that benefits from high-content microscopy approaches like SPI. Notably, SPI generates instant sub-diffraction-limited images with a line readout rate exceeding 10 kHz (92,500 μm^2^/s, **Supplementary Videos S5-6**) without degradation across milli-meter-scale FOV (**Fig. 3c, Supplementary Videos S7-10**). This high scalability contrasts with traditional wide-field microscopy, which cannot simultaneously capture both subcellular details and macroscopic multicellular structures (**Fig. S6**). The SPI results revealed key adaptive phenotypes that evolved over 1,000 days (~5,000 generations, **Fig. 3d**), including progressive cellular elongation with average aspect ratios increasing from ~1.30 (ancestors) to ~3.28 by 1,000 transfers (all samples taken from anaerobic population PA5; **Fig. 3e**). These morphological changes transformed module architecture. While the ancestor formed clusters with a mean size of 30.7 µm, these modules increased to 87.1 µm after 1000 transfers (**Fig. 3f**). Interestingly, module size is approximately scaled linearly with cellular aspect ratio during early evolution, but this relationship diverged at two key timepoints (t400 and t1000; **Fig. 3g**), hinting at the co-evolution of adaptations that increase module size by reducing within-module strain accumulation or increasing mechanical toughness. This evolutionary trajectory illustrates how simple multicellular groups can rapidly evolve increased size and material robustness through innovations in cellular morphology and attachment patterns. SPI can play an important role in evolutionary cell biology, providing deep datasets characteristic of widefield microscopy while retaining sub-cellular precision.

## DISCUSSIONS

In conclusion, the SPI system addresses current demands in large-scale super-resolution cell analysis by enabling resolution doubling, continuous and steady throughput, unconstrained FOV, and instantaneous super-resolution image creation, notably at a much simpler configuration compared to pioneering works^16,32–35^ (**Figure S14**). This approach features highly desirable adaptability in instrumentation, protocols, and scalable cellular environments^36^, whose functionality can be further extended with optical and computational strategies^37–41^. For example, the high sensitivity of SPI allows for probing living cells through dynamic endogenous labels such as autofluorescence (**Fig. S7, Supplementary Video S11**). In addition, non-iterative rapid WB deconvolution can be replaceable using unsupervised networks^42^ for greater computational flexibility (**Fig. S8**). Furthermore, the epi-fluorescence platform presents great potential for integration with high-throughput screening^43^, imaging flow cytometry^44^, fluorescence lifetime imaging^39^, and spatial-resolved transcriptomics^45,46^. We anticipate that the SPI technique will provide a methodological pathway for elucidating fundamental and translational biological systems beyond the optical and computational limits.

## METHODS

### Super-resolution imaging system

The system was constructed using an epi-fluorescence microscope (Nikon Eclipse Ti2-U) equipped with a 100×, 1.45 NA oil-immersion objective lens (Nikon CFI Plan Apochromat Lambda 100× Oil), following established configurations^19,20,47^. For excitation, a 488-nm laser (Coherent OBIS LX) generated multifocal illumination, which was spatially modulated by a microlens array (MLA; RPC Photonics MLA-s100-f4-A-R1; 100-µm pitch, F-number 4.2). The MLA produced a grid of focal points demagnified to a 1.6-µm pitch at the sample plane via relay optics. Since the demagnified diffraction limit of the MLA is much smaller than the objective’s diffraction limit, the chosen F-number of 4.2 ensures that each excitation focus remains diffraction-limited by the objective lens at the sample plane (see **Supplementary Note 1**). Fluorescent emissions were filtered using a dichroic mirror (Chroma ZT 405/488/561/647 RPC-UF1) and emission filter (Chroma ZET 405/488/561/647 m) to isolate target signals. For detection, a custom MLA (160-µm pitch, F-number 2.1) was positioned at the image plane. The MLA was fabricated via two-photon 3D lithography (Nanoscribe GT2, 10× objective lens, IP-n162 photoresist) on the fused silica substrate at the Institute for Electronics and Nanotechnology at Georgia Institute of Technology. The MLA enabled optical photon reassignment and was mounted with three-axis adjustability (x, y, rotation) to align precisely with the emission grid coordinates. Reassigned signals were captured by a timedelay integration (TDI) camera (Vieworks VT-4K5X-H200; 5-µm pixel size) paired with a macro lens (Nikon AF-S VR Micro-NIKKOR; 105 mm, f/2.8G IF-ED; forming 1.44× demagnification). The effective magnification of the optical system is approximately 69×, yielding approximately ~72 nm per pixel. Full component specifications are provided in **Table S2**.

### Image acquisition and reconstruction

After aligning the system, the stage controller (ASI MS-2000-500) was set to move at a speed corresponding to the line rate of the TDI sensor at the sample plane. A series of 12-bit depth images were acquired, with each image seamlessly appended to form a panoramic view according to the predefined dimensions of the buffer height. Here, surface imperfections from the fabrication of the microlens array introduced inherent fixed-pattern noise parallel to the scan direction. As shown in **Supplementary Note 5**, a fluorescent calibration slide was used to measure the time-averaged intensity variation of the accumulated line and effectively remove the fixed-pattern noise. Forming a large FOV through raster scanning further requires stitching of multiple panoramically reconstructed scans^48^. Due to the uneven signalto-noise ratio (SNR) across each scanned image, which is higher toward the center, large FOV images may exhibit periodic fluctuations. These artifacts can be mitigated by applying notch filtering, using finer scanning steps to ensure sufficient SNR, or increasing the beam expansion ratio for more uniform illumination.

Achieving a full twofold resolution improvement requires an additional step of WB deconvolution. The theoretical 2× improved resolution of green fluorescence is approximately 110 nm, whereas an instant intermediate image with a 72-nm pixel size does not meet the criteria for proper sampling. Thus, the instant image is upscaled into 36 nm per pixel with bilinear interpolation and deconvolved. For deconvolution, we assume a theoretical PSF modeled as a 2D Gaussian distribution with a wavelength of approximately 370 nm, reduced by a factor of √2 from 520 nm, since the Rayleigh resolution scales linearly with wavelength.

### HeLa cell culture and immune labeling

HeLa cells (Sigma-Aldrich, #93021013) were cultured in Dulbecco’s modified Eagle medium (DMEM) with 10% fetal bovine serum (FBS) and 1% Penicillin-Streptomycin (Pen-Strep) at 37 °C and in a 5% CO_2_ atmosphere.

On the day of immune labeling microtubule, the cells were fixed with 0.3% (volume:volume) glutaraldehyde in extraction buffer, incubating for 1 minute at 37 °C. The extraction buffer consists of 10 mM MES, 150 mM NaCl, 5 mM EDTA, 5 mM glucose, 5 mM glucose, 5 mM MgCl2, and 0.25% (volume:volume) Triton X-100 in ultra-pure water. The buffer of the cells is switched to 2% (volume:volume) glutaraldehyde in cytoskeleton buffer at room temperature for 10 minutes. The cytoskeleton buffer is the aforementioned extraction buffer without the Triton X-100. The cells are then gently washed with blocking/permeability (b/p) solution for 5 minutes, 3 times. The b/p solution consists of 2.5% (weight:volume) bovine serum albumin (BSA) and 0.1% (volume:volume) Triton X-100 in phosphate-buffered saline (PBS, Corning, #21-040-CM). Then, the primary antibody that targets β-tubulin was added to the cell dish with 2 mL of b/p solution at a concentration of 2 µg/mL (ThermoFisher, #32-2600). The cell dish was placed inside a humidified chamber at room temperature for 1 hour. After primary antibody tagging, the cell dish was washed with b/p solution 3 times, 5 minutes each time. Then, the cells were labeled with 2 µg/mL of Goat anti-mouse IgG conjugated with Alexa Fluor Plus 488 (ThermoFisher, #A32723). The staining occurred inside the humidified chamber at room temperature for 1 hour. After secondary antibody labeling, the cell dish was washed with b/p solution three times 5 minutes each and sequentially washed with PBS twice. The sample is finally stored at 2 mL of PBS solution for imaging.

### U2OS cell culture and mitochondria staining

U2OS cells (Sigma-Aldrich, #92022711) were cultured in DMEM with 10% FBS and 1% Pen-Strep at 37 °C and in a 5% CO_2_ atmosphere. A day before imaging, the cells were incubated with a pre-warmed (37 °C) mixed solution containing 3 mL modified DMEM and 20 µL CellLight Mitochondria-GFP (ThermoFisher, #C10600). The GFP was expressed on the mitochondria after 18 hours of incubation. On the day of imaging, the growth medium was removed, and the cells were fixed with 4% paraformaldehyde (PFA, diluted from 16% PFA with PFA:PBS:ultrapure-water with a 1:2:1 ratio, Electron Microscopy Sciences) at room temperature for 12 minutes. The cells were washed with a clear PBS solution for imaging.

### Fluorescence staining of peripheral blood smear

Peripheral blood smears were prepared following a standard protocol^27^. A sterile lancet was used to obtain a drop of capillary blood from the fingertip, which was immediately placed on a #1.5 coverslip (VWR, #48366). The drop was spread using a glass slide at a 30° angle to create a thin, mono-layered smear. The slides were air-dried for 5 minutes and subsequently fixed with 1 mL HPLC methanol (Sigma-Aldrich, #34885) for 10 minutes at room temperature. Following the fixation, the slides were airdried for 10 minutes and subsequently stained with Alexa Fluor 488 conjugate wheat germ agglutinin (ThermoFisher, #W11261) at 2% (volume:volume) dilution in PBS for 15 minutes at room temperature. After staining, the slides were gently washed with PBS to remove excess dye and allowed to air dry. Finally, a drop of SlowFade Gold Antifade Mountant (ThermoFisher, #S36936) was applied before placing it on the glass slide for imaging.

### Population analysis

The population of each cell type in the peripheral blood smear was quantified from images obtained using the SPI system. For erythrocytes and platelets, total counts were extrapolated from multiple small inspection areas, averaging across several measurements to ensure accuracy. Erythrocytes were estimated at approximately 82,000 ± 6,500, and platelets at around 5,600 ± 660 (mean ± standard error). Leukocytes were manually counted across the entire inspected area, using prior knowledge of cell size and nuclear morphology to distinguish different subtypes.

### Snowflake yeast

Snowflake yeast (*Saccharomyces cerevisiae*) was derived from an engineered diploid Y55 strain, in which both copies of the *ACE2* gene were deleted, leading to incomplete cell separation and stable cluster formation. Obligate anaerobic ancestors were established by selecting a spontaneous petite mutant characterized by mitochondrial DNA deletions that prevent respiration. Yeast cultures were maintained in YPD medium (1% yeast extract, 2% peptone, 2% dextrose) at 30 °C with shaking at 225 rpm. For the multicellularity long-term evolution experiment (MuLTEE)^29,30^, anaerobic populations (PA) underwent daily settling selection to promote the evolution of larger clusters. Cultures were grown in 10 mL of YPD medium for 24 hours before selection. For each round of selection, 1.5 mL of culture was transferred to an Eppendorf tube, left undisturbed for 3 minutes, and the top 1.45 mL was removed, retaining only 50µL of the settled fraction for reinoculation. As clusters increased in size, the settling time was gradually reduced to 30 seconds after ~350-500 generations.

We used genetically modified strains from the anaerobic lines where the eGFP tag is inserted into the genome downstream of a copy of the PDC-1 gene in the ancestor and the HSC-82 gene in the ancestor, 400, and 1000-day evolved strains. eGFP tagging was done using homologous recombination. We also used 400 and 600-day evolved strains with SS-GFP sequence^49^ under a strong TPI1 promoter at the HO locus to visualize the endoplasmic reticulum. To prepare samples for imaging, frozen cultures of the aforementioned GFP-tagged strains were thawed and streaked on YPD-agar (1% agar + YPD) plates. Liquid cultures were inoculated from single colonies in YPD and grown for three days with settling selection to reach equilibrium. A 1 mL sample was collected, and the YPD medium was replaced with PBS by centrifuging at 5000 g for 1 minute, followed by aspiration. This process was repeated twice to minimize autofluorescence from the culture medium. The collected clusters were evenly spread onto a positive-charged glass slide and gently fragmented into smaller modules by a coverslip before imaging.

### Measuring cell aspect ratio and cluster module size

The aspect ratio of individual cells was calculated from images of cells obtained with the SPI system. Around 50 individual cells per evolutionary stage were collected for measurement using ImageJ. To measure the size of multicellular groups, we used ImageJ to threshold and segment SPI images of snowflake yeasts, which allowed us to measure the cross-sectional area of individual clusters of around 20-30 modules per evolutionary stage.

## DATA AVAILABILITY

Data underlying the results presented in this paper may be obtained from the corresponding author upon reasonable request.

## CODE AVAILABILITY

The code is written in MATLAB (tested in 2020b, MathWorks). The latest version of the software is available at https://github.com/ShuJiaLab/SPI

## ACKNOWLEDGMENTS

This work is supported by the Parker H. Petit Institute for Bioengineering and Biosciences of Georgia Institute of Technology, NSF grants EFMA1830941 (to S.J.) and DBI2145235 (to S.J.), and NIH grants R35GM124846 (to S.J.) and R35-GM138030 (to W.C.R). This work was performed in part at the Georgia Tech Institute for Electronics and Nanotechnology, a member of the NSF-supported National Nanotechnology Coordinated Infrastructure (ECCS-1542174).

We thank Dr. Parvin Forghani Esfahani of Dr. Chunhui Xu’s laboratory at Emory University for kindly providing human-induced pluripotent stem cell (hiPSC)-derived cardiac fibroblast samples used in this study. We also thank Nikolas Roeske at the Georgia Tech Institute for Electronics and Nanotechnology for his instruction and training in Nanoscribe 3D lithography, which enabled the fabrication of the custom microlens arrays.

## AUTHOR CONTRIBUTIONS STATEMENT

K.Y., B.M., and S.J. conceived and designed the project. K.Y., B.M., and K.T. contributed to the construction of the optical system. K.Y. and H.Y. performed imaging experiments. K.Y. and H.Y. conducted image processing. K.Y., H.Y., K.T., Z.L., S.D., G.O.B., and A.J.B. prepared biological samples. W.C.R. provided biological insights. S.J. supervised the overall project. K.Y. and S.J. wrote the manuscript with input from all authors.

## COMPETING INTERESTS STATEMENT

S.J., K.Y., and H.Y. are listed as inventors on a pending patent application (US Patent App. 63/814,286), partly relevant to the technology described in this manuscript. The other authors declare no competing interests.

